# Electric Field-Induced Destabilization and Surface Modulation of Aβ42 Fibrils in Molecular Simulations: Theoretical Implications for DC Stimulation in Alzheimer’s Disease

**DOI:** 10.1101/2025.09.24.678267

**Authors:** Fran Bačić Toplek, Natale Vincenzo Maiorana, Matteo Guidetti, Sara Marceglia, Riccardo Capelli, Alberto Priori, Carlo Camilloni

## Abstract

The amyloid-β peptide 42 (Aβ42) forms fibrillar aggregates that are a hallmark of Alzheimer’s disease. While recent pharmacologic therapeutic strategies targeting Aβ42 fibrils and oligomers have shown promising results, safer and more effective approaches are still needed. Non-invasive brain stimulation (NIBS) techniques such as repetitive transcranial magnetic stimulation (rTMS) and transcranial direct current stimulation (tDCS) have been increasingly explored as a possible complementary intervention, but the molecular mechanisms by which static electric fields could influence amyloid aggregation remain poorly understood. Here, we use atomistic molecular dynamics simulations to investigate the effects of static electric fields on Aβ42 fibrils. We examine the response of an ex vivo fibril structure, with reconstructed N-terminal regions, to increasing field strengths under different structural restraint conditions. Our results show that the electric field perturbs the disordered N-terminal “fuzzy coat,” altering its conformational dynamics and reducing its interactions with the fibril core. This reorganization modifies the surface properties of the fibril, potentially impairing secondary nucleation. Additionally, simulations with unrestrained fibril ends reveal increased fluctuations in core residues, particularly near the N-terminus, indicating a destabilizing effect that may hinder fibril elongation. While the field strengths used here exceed those typical of NIBS, our findings provide a molecular-level rationale for how electric fields could modulate fibril propagation and support further experimental investigations under physiologically relevant conditions.

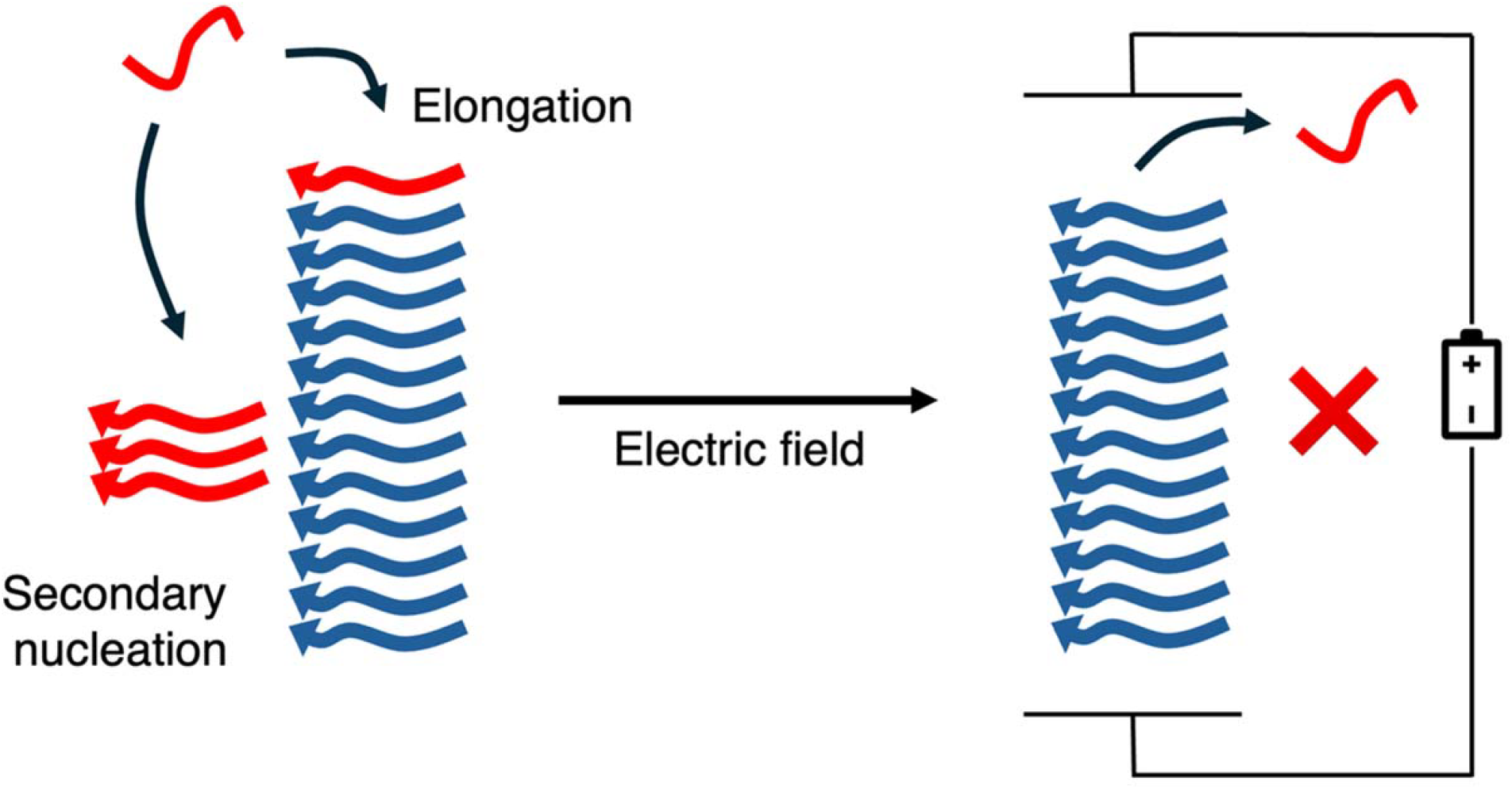

## Introduction

The amyloid-β peptide 1-42 (Aβ42) is the main protein component of extracellular plaques found in the brains of Alzheimer’s disease (AD) patients^1,2^. Fibrils of wild-type Aβ42, isolated from postmortem AD brain tissue, have been resolved by cryo-electron microscopy and shown to consist of two identical S-shaped protofilaments embracing each other with extended arms. The protofilament core is made by residues 9 to 42, flanked by a disordered, solvent-exposed coating constituted by the N-terminal residues 1–8^3,4^, as illustrated in **Figure 1**. *In situ* cryo-electron tomography has further revealed that β-amyloid plaques contain a parallel array of fibrils organized into lattice-like assemblies^5^.

**Figure 1:**
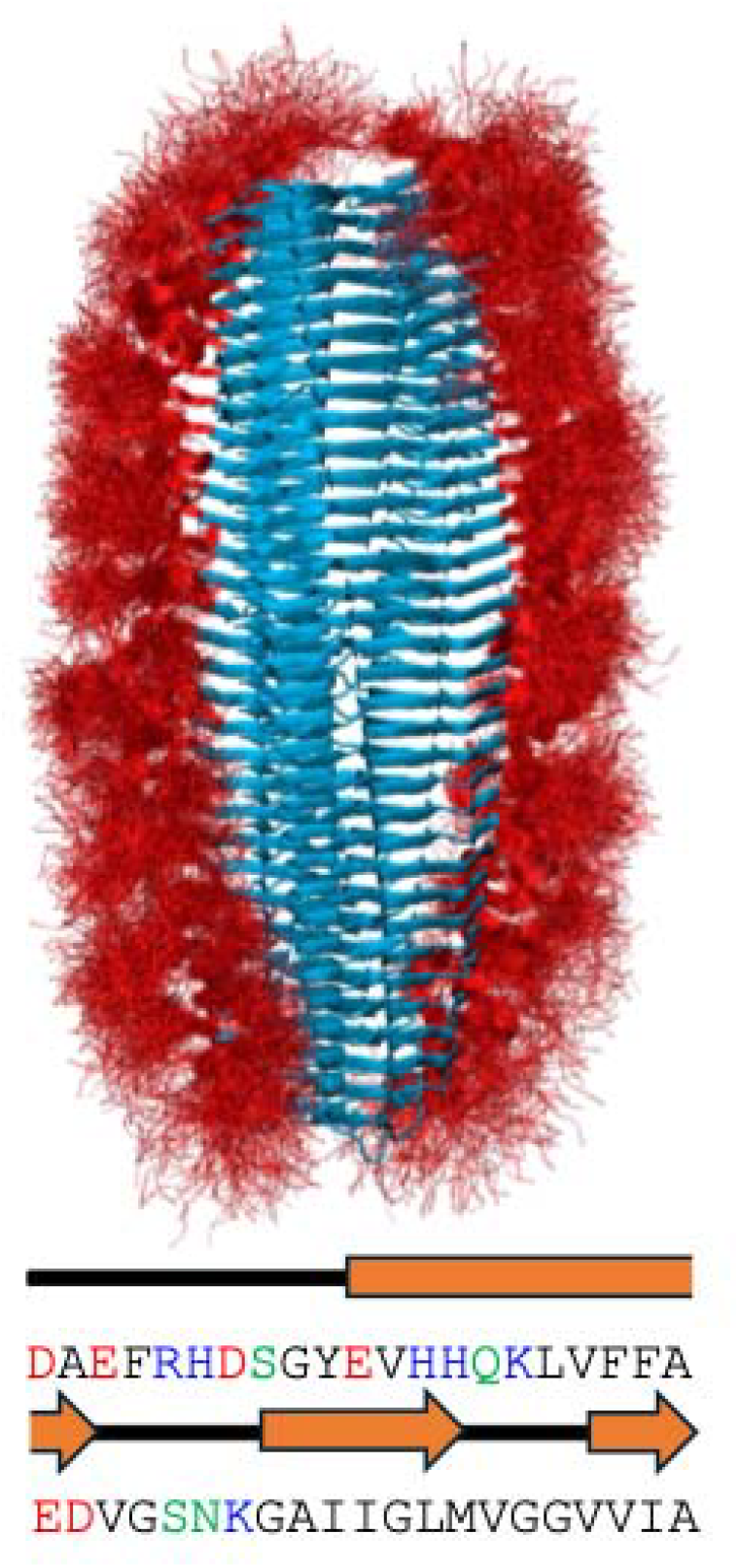
Representation of the structure of Aβ42 simulated without electric field (EF). The fibril core region is represented in cartoon highlighting the parallel beta structure while the N-terminal tails are shown as ribbons. At the bottom is reported the protein sequence and secondary structure as observed in the fibril, amino acids are colored by their physicochemical classification as acidic (red), basic (blue), polar (green), and hydrophobic (black).

*In vitro* studies of Aβ42 aggregation have uncovered a complex self-assembly mechanism, involving a combination of primary nucleation and surface-catalyzed secondary nucleation processes, where the latter is deemed to be responsible to produce cytotoxic oligomeric species^6–8^. The monomeric Aβ42 peptide, produced by proteolytic cleavage of the amyloid precursor protein, is intrinsically disordered and samples a broad range of conformations on the microsecond timescale^9–11^. Systematic mutational analyses have shown that hydrophobic or negatively charged variants in the polar segment of the sequence (residues 1–28, cf. **Figure 1**) generally reduce secondary nucleation, whereas most other mutations in that region tend to enhance it. In contrast, mutations in the apolar C-terminal region (residues 29–42) more frequently reduce secondary nucleation^12–15^. These findings support a key role for the disordered N-terminal region in promoting fibril proliferation via the recruitment of monomers in secondary nucleation. Such mechanistic insights are relevant for therapeutic development, as interventions targeting fibril proliferation represent a promising strategy to limit disease progression.

Recent breakthroughs in AD pharmacotherapy have demonstrated that antibodies targeting Aβ fibrils and oligomers can slow disease progression^16^. Nevertheless, there remains a need for more effective and potentially complementary therapeutic strategies. In this context, non-invasive brain stimulation (NIBS) techniques like repetitive transcranial magnetic stimulation (rTMS) and transcranial direct current stimulation (tDCS) have emerged as a promising approach to alleviate AD symptoms^17^ and enhance neuronal plasticity^18^. Furthermore, they were suggested to possibly facilitate Aβ clearance by transiently increasing brain-blood barrier permeability^19,20^, Aβ transport^21^, and microglial phagocytosis of Aβ^22^. However, the molecular effects of electric fields (EFs) on protein aggregation remain poorly understood. A recent molecular dynamics (MD) study by Kalita et al.^23^ reported that static and oscillating (with frequency in the GHz range) electric fields applied to aggregates of small portions of Aβ42 prevent their formation and contribute to their disassembly. The same group and others also observed in simulations the disruptive effect of electric fields on trimers composed by full Aβ42 peptides^24^ and other small oligomers^25^. Similarly, simulations of few layers of α-synuclein fibrils under static electric fields suggested the possibility of a progressive break of β structures, causing destabilization of the fibril core after an EF threshold value of approximately 0.3–0.4 V/nm^26,27^. However, it remains unclear whether these observations extend to full-length Aβ42 fibrils and whether EFs can perturb specific regions of biological relevance such as the flexible N-terminal tails or the elongation surfaces.

Motivated by in-tissue observations of highly ordered fibril arrangements, in this work we investigate the molecular impact of a static electric field aligned with the fibril axis on a pre-formed Aβ42 fibrils. We focus on two key aspects: the conformational dynamics of the disordered N-terminal tails (connected with secondary nucleation) and the structural stability of the fibril’s top layer (involved with the elongation of the fibril). Our results suggest that the electric field may reduce secondary nucleation by perturbing the conformational dynamics of the N-terminal fuzzy region and modifying the properties of the fibril surface. In parallel, the field appears to promote progressive destabilization and dissolution of fibril ends involved in elongation.

## Results and Discussion

We performed MD simulations of an Aβ42 fibril composed of two protofilaments, each containing 33 monomers, after modeling the missing N-terminal residues (residues 1–8). Simulations were carried out in triplicate across a range of static EF strengths (0 to 0.2 V/nm), and two restraint conditions were considered: (i) positional restraints on the entire fibril core, and (ii) positional restraints excluding the top two layers of the core (see *Materials and Methods* for details). The rationale behind this choice was to mitigate finite-size effects that could arise due to the limited length of the simulated fibril. Each simulation was run for 1 to 2 μs, and analysis was performed after discarding the first 500 ns to allow for initial relaxation of the system.

### A static electric field perturbs the disordered N-terminal region

To investigate the effect of a static EF on the disordered N-terminal “fuzzy coat”, we first analyzed simulations in which the fibril core was held fixed by positional restraints on the Cα atoms. This setup preserved the flexibility of the N-terminal tails while minimizing structural rearrangements in the ordered core.

**Figure 2A** shows the root mean square deviation (RMSD) of the average intermolecular distances between residues in the fuzzy coat, used here as a measure of their relative displacement under the applied field. This displacement increases with field strength and saturates above ∼0.08 V/nm. Visual inspection reveals that the N-terminal tails, which carry a net charge of -3, tend to align with the direction of the field.

**Figure 2:**
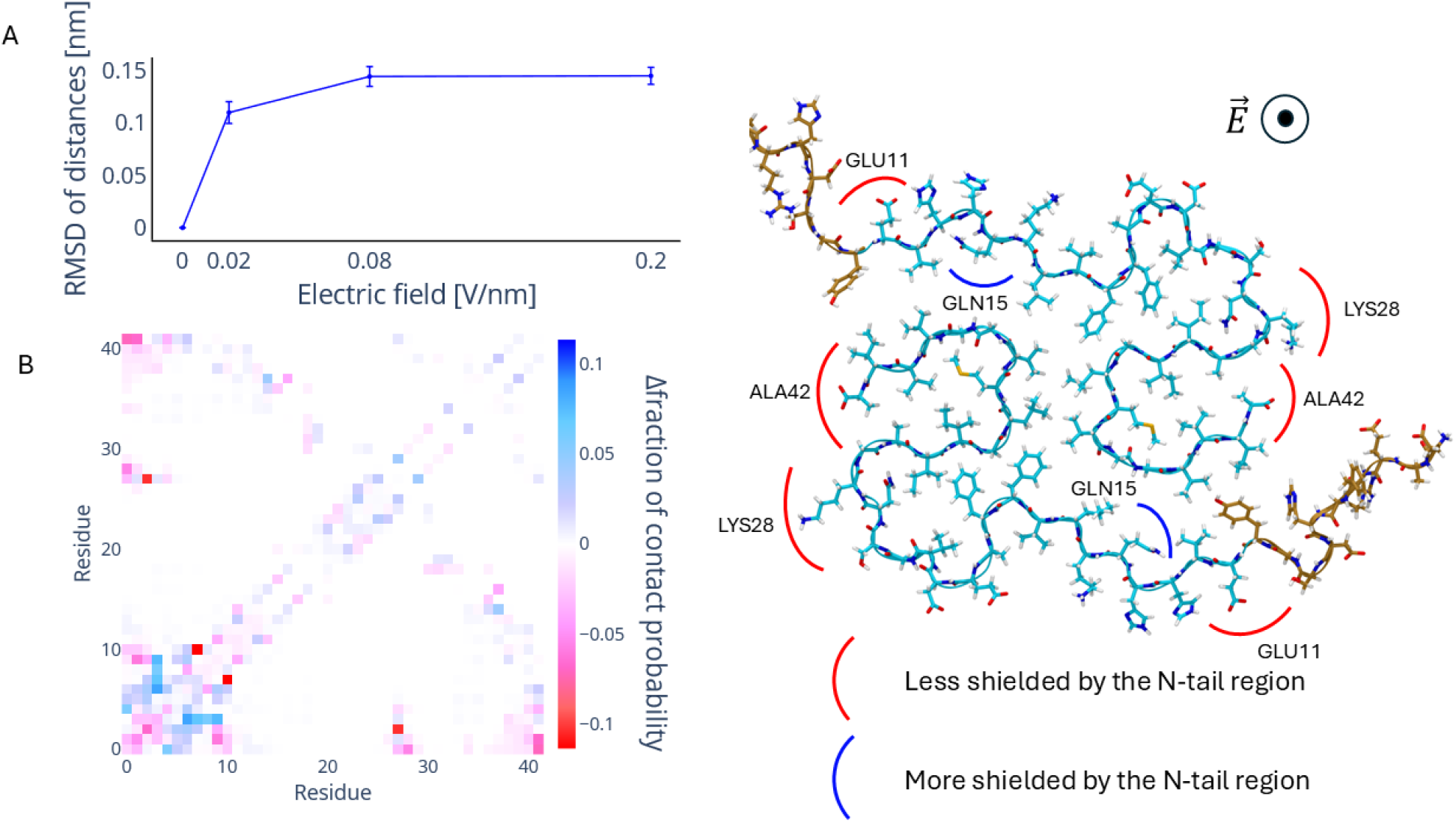
A) The root-mean-square deviation (RMSD) of average inter-residue distances over the fibril as a function of the electric field strength. The RMSD increases until saturation is reached at around 0.08 V/nm. B) A contact map showing the difference in contact formation probability given the intermolecular residue pairs in the fibril. The difference is calculated from the contact map of the fibril in the 0.08 V/nm electric field and the reference without an electric field applied. C) The structure of a layer of the fibril, residues losing (less shielded) or gaining (more shielded) a significant fraction of contacts are highlighted.

To assess how this displacement affects interactions with the fibril surface, we computed the intermolecular residue–residue contact probability, defined as the fraction of frames in which any two atoms from distinct residues are within 0.5 nm. **Figure 2B** shows the difference in contact probability matrices between the 0.08 V/nm and zero-field simulations. A systematic loss of contacts is observed between the N-terminal tails and the fibril core, as well as among the tails themselves. This effect is particularly pronounced for several surface core residues, including E11, K28, and A42 as shown in **Figure 2C**. Only Q15 forms more contacts under the effect of the EF. This reorganization likely alters the physico-chemical properties of the fibril surface, leading to a reduction in its hydrophobic character. We hypothesize that this change may impair the catalytic activity of the fibril surface in secondary nucleation processes.

### A static electric field destabilizes the fibril core ends

To assess the effect of EFs on fibril stability, we performed additional MD simulations in which the top two monomer layers were left unrestrained, while the remainder of the fibril core was restrained as in the previous setup. This configuration allowed us to specifically probe the impact of the EF on the structural stability of the fibril ends—key regions for fibril elongation— while mitigating finite-size artifacts associated with the relatively short fibril model. Results of these simulations are illustrated in **Figure 3**, which shows the RMSD of the two top layers of core monomers over time and a representative final conformation.

**Figure 3:**
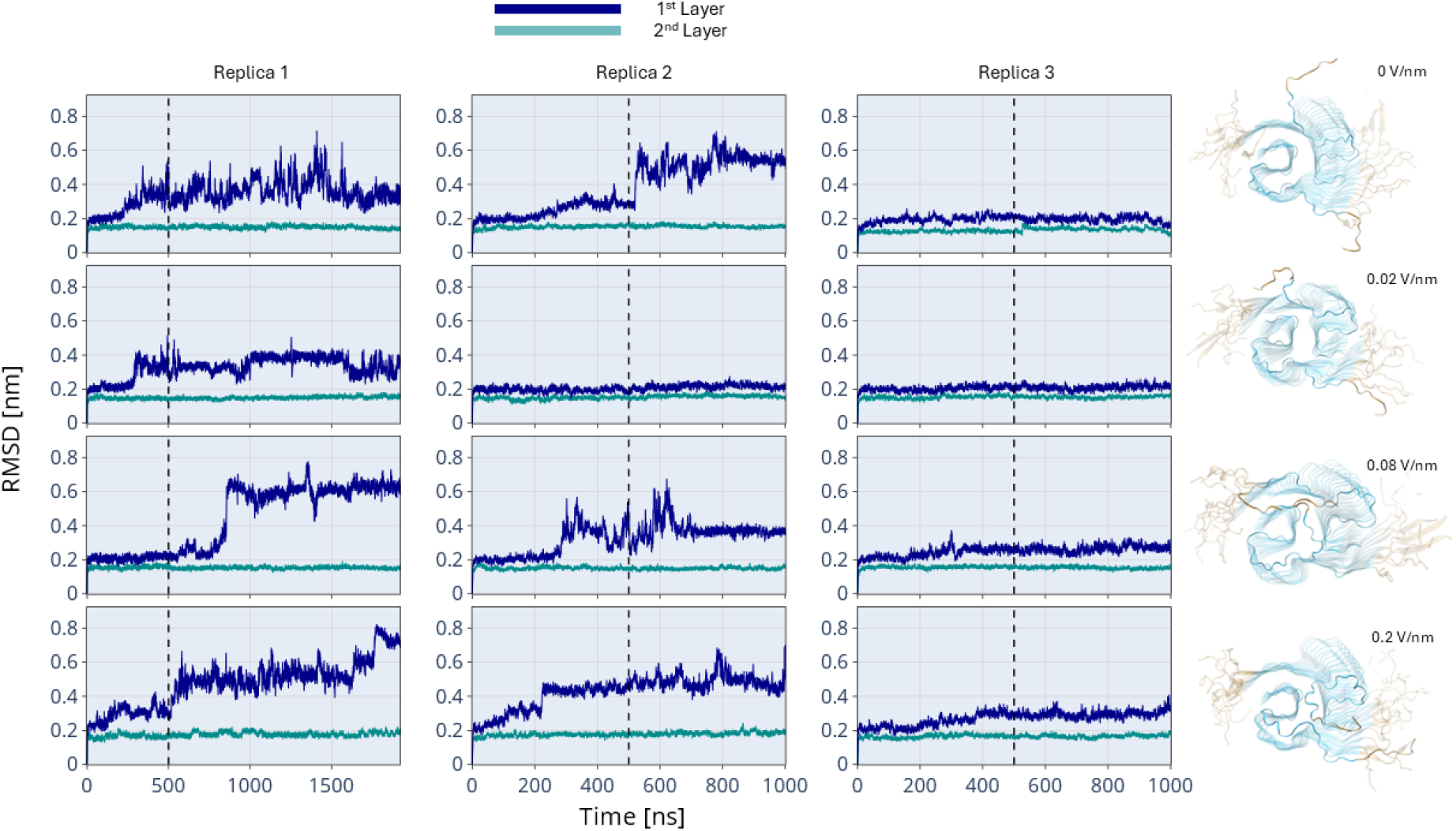
Effect of EF intensity on the stability of the top two fibril monomers. In the first three columns, the RMSD of the two unrestrained monomers from the upper layer (blue) and the adjacent layer below (cyan) is shown together for each EF intensity and simulated replica, relative to the minimized starting structure. EF intensities are arranged row-wise and replicas column-wise. The last column shows a representative fibril conformation after MD, with the top layer in solid representation.

On the microsecond timescale accessible to our simulations and within the range of EF strengths employed, we did not observe complete monomer dissociation. However, we detected qualitative structural perturbations in the unrestrained regions of the core of the top layer irrespectively of the EF intensity employed. On the contrary the second top layer seems stable in all simulations. Such results suggest that focusing on the second layer is more reliable as conformational dynamics of the first is too noisy.

To localize the destabilizing effect more precisely, we analyzed the root-mean-square-fluctuations of atomic positions (RMSF) between identical residues of the second unrestrained fibril layer. For each residue, we computed the RMSF and compared it between simulations with and without the EF. An increase in RMSF indicates enhanced fluctuation, which we interpret as a reduction in local stability.

**Figure 4A** reports the residue-wise difference RMSF at 0.08 V/nm relative to the zero-field case. The results show a marked increase in fluctuations for core residues 11, 12, 15, and 23, suggesting that the increased mobility of the disordered N-terminal region under the field propagates into the structured fibril core electric fields affects individual charged residues further downstream. Notably, we observe no significant decrease in fluctuations at any residue position, indicating that the destabilizing effect of the electric field is both localized and systematic.

**Figure 4:**
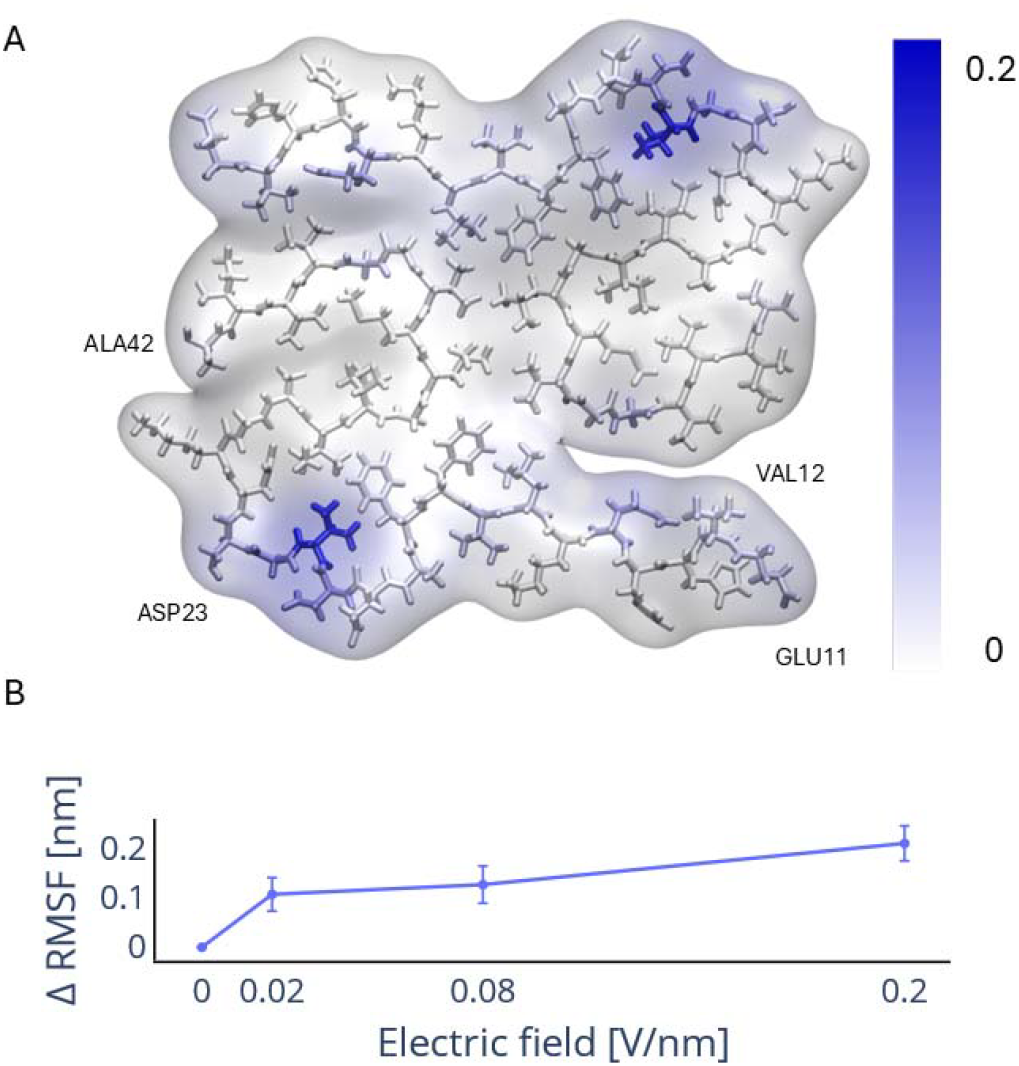
A) Surface and licorice representation of the second unrestrained layer monomers in the Aβ42 fibril colored by their change of RMSF for an electric field of 0.08 V/nm. B) RMSF as a function of the electric field strength.

Averaging the residue-wise RMSF as a function of EF strength (see **Figure 4B**) we found a trend similar to that observed for the displacement of the N-terminal tails, i.e., fluctuations in the fibril core increase with field strength, but do not appear to saturate. Instead, the fluctuations keep increasing with the electric field.

### Mechanistic hypothesis and study limitations

Together, our results suggest a twofold mechanism by which a static EF perturbs Aβ42 fibril structure and dynamics. First, the field reorients and displace the disordered N-terminal tails, which are rich in charged residues, leading to a substantial changes to the fibril core surface. This effect likely modifies the surface properties that are key to attract new monomers, potentially decreasing its catalytic activity in surface-mediated secondary nucleation. Second, the field appears to destabilize the fibril ends—particularly residues 11–15, and 23—by enhancing local structural fluctuations, which may impair elongation. These observations, together with those recently published for small oligomers of Aβ and very small ⍰-synuclein fibrils support the hypothesis that an external EF could modulate fibril propagation by interfering with both primary and secondary nucleation as well as elongation^23–28^.

These molecular insights come with important limitations. Most notably, the EF strengths employed in this study—as in previous computational works—are several orders of magnitude higher than those estimated to arise from tDCS^29^. This choice is necessary to observe conformational changes within the accessible microsecond simulation timescale. Nevertheless, we can foresee that weaker fields could produce similar effects over longer, physiologically relevant timescales. In our simulations, we applied a constant, spatially uniform electric field aligned with the fibril axis, whereas in vivo conditions are likely to involve heterogeneous fibril arrangements and time-dependent field orientations. We anticipate that fields in virtually any direction would disrupt the conformational dynamics of the highly charged, disordered N-terminal region, which supports our hypothesis that secondary nucleation processes are perturbed. In contrast, the destabilization of the fibril core that we observed appears to be a specific consequence of fields oriented along the fibril axis.

In conclusion, our findings provide a molecular-level rationale for how static electric fields can modulate amyloid fibril behavior by perturbing both the dynamics of the disordered surface and the stability of the fibril end. These results deepen the mechanistic understanding of fibril propagation under external electric fields and offer molecular support for their potential therapeutic use. We encourage experimental efforts to test these predictions by measuring Aβ42 aggregation kinetics in vitro under static EF conditions.

## Material and Methods

All simulations were performed using GROMACS 2024.4^30^ on the LEONARDO supercomputer at CINECA. The DES-AMBER force field ^31^ was used in combination with the TIP4P-D water model^32^ and a 200 mM ionic concentration to neutralize the system. The system contained 95,350 molecules of water, 398 sodium ions, 200 chloride ions and 66 Aβ42 monomers.

The starting fibril structure was derived from PDB entry 7Q4B^3^ and extended to 33 layers, each comprising two monomers, following the reported fibril symmetry. The full construct included intrinsically disordered N-terminal regions, introduced by Modeller^33^, amounting to the first 8 residues, which were added to all monomers. The system was solvated in a rectangular box of dimensions 13.11 × 12.69 × 19.79 nm^3^.

Electric fields along the z-axis between 0 and 0.2 V/nm were treated as static, based on the assumption that alternating fields would appear quasi-static on the microsecond timescale relevant for the simulations. All systems were initially energy-minimized by steepest descent to a maximum force of 200. The c-rescaling algorithm^34^ and v-rescaling algorithm^35^ were used to maintain constant pressure of 1 atm (coupling time 5 ps) and temperature of 310 K (coupling time 1 ps) during production runs. Electrostatic interactions were handled using the Particle Mesh Ewald method^36^ with a short-range cutoff of 1.0 nm and a Fourier grid spacing of 0.12 nm. Van der Waals interactions were treated with a 1.0 nm cutoff. Covalently bonded hydrogens were constrained using the LINCS algorithm^37^.

Two types of simulation setups were used. In the core-restrained simulations, positional restraints were applied to Cα atoms of residues 11–42 in all layers, allowing N-terminal flexibility while maintaining the core structure. In the top-layer-unrestrained simulations, only the four topmost monomers were left unrestrained, with restraints applied to all other core residues (11– 42), enabling potential detachment or restructuring under electric field influence.

Each simulation was run in three replicates with one replica ran for 2 μs and two for 1 μs. The first 500 ns were discarded from analysis to allow for relaxation of the fibril structure. The full simulation trajectories can be found in the Zenodo database under the DOIs: 10.5281/zenodo.17160960 and 10.5281/zenodo.17161058.

## AUTHOR INFORMATION

### Author Contributions

The manuscript was written through contributions of all authors. All authors have given approval to the final version of the manuscript.

## ACKNOWLEDGMENT

The authors acknowledge CINECA for an award under the ISCRA initiative, for the availability of high-performance computing resources and support.

## ABBREVIATIONS

NIBS: Non-invasive brain stimulation
rTMS: repetitive transcranial magnetic stimulation
tDCS: transcranial direct current stimulation
EF: Electric field
MD: molecular dynamics
RMSD: root mean square deviation
RMSF: root mean square fluctuations

